# Human cytomegalovirus vMIA inhibits MAVS oligomerization at peroxisomes in an MFF-dependent manner

**DOI:** 10.1101/2022.02.25.481859

**Authors:** Ana Rita Ferreira, Ana Gouveia, Ana Cristina Magalhães, Isabel Valença, Mariana Marques, Jonathan C. Kagan, Daniela Ribeiro

## Abstract

Upon intracellular recognition of viral RNA, RIG-I-like proteins interact with MAVS at peroxisomes and mitochondria, inducing its oligomerization and the downstream production of direct antiviral effectors. The human cytomegalovirus (HCMV) is able to specifically evade this antiviral response, via its antiapoptotic protein vMIA. Besides suppressing the programmed cell death of infected cells, vMIA inhibits the antiviral signalling at mitochondria by inducing the organelle’s fragmentation, consequently hindering the interaction between MAVS and the endoplasmic reticulum protein STING. Here, we demonstrate that vMIA interferes with the peroxisomal antiviral signalling via a distinct mechanism that is independent of the organelle’s morphology and does not affect STING. vMIA interacts with MAVS at peroxisomes and inhibits its oligomerization, restraining downstream signalling, in an MFF-dependent manner. This study also demonstrates that vMIA is totally dependent on the organelle’s fission machinery to induce peroxisomal fragmentation, while this dependency is not observed at mitochondria. Furthermore, although we demonstrate that vMIA is also able to inhibit MAVS oligomerization at mitochondria, our results indicate that this process, such as the whole vMIA-mediated inhibition of the mitochondrial antiviral response, is independent of MFF. These observed differences in the mechanisms of action of vMIA towards both organelles, likely reflect their intrinsic differences and roles throughout the viral infection.

This study uncovers specific molecular mechanisms that may be further explored as targets for antiviral therapy and highlights the relevance of peroxisomes as platforms for antiviral signalling against HCMV.

## Introduction

Upon viral infection, the intracellular retinoic acid-inducible gene-I (RIG-I)-like receptors (RLRs), such as RIG-I and melanoma differentiation-associated gene-5 (MDA-5), interact with viral RNA (Saito and Gale, 2008) and subsequently activate the adaptor protein mitochondrial antiviral signalling (MAVS) at mitochondria (Kawai et al., 2005; Meylan et al., 2005; Seth et al., 2005; Xu et al., 2005), peroxisomes (Dixit et al., 2010) and mitochondrial-associated membranes (MAMs) (Horner et al., 2011). Interaction between the caspase activation and recruitment domains (CARDs) of both RLR and MAVS, induces MAVS oligomerization and amplifies antiviral signalling (Hou et al., 2011; Seth et al., 2005), culminating with the production of interferons (IFNs) and IFN-stimulated genes (ISGs) (Dixit and Kagan, 2013; Kell and Gale, 2015). Together, peroxisomes and mitochondria orchestrate the antiviral immune response mediated by MAVS: peroxisomal MAVS leads to a rapid expression of ISGs, which is then complemented by the mitochondrial counterpart, prompting a long-term, more stable and amplified response (Dixit et al., 2010). The importance of peroxisomes for the cellular antiviral response is highlighted by recent studies demonstrating that distinct viruses, such as the human cytomegalovirus (HCMV) (Magalhães et al., 2016; Marques et al., 2018), dengue and West Nile viruses (You et al., 2015), hepatitis C virus (Bender et al., 2015; Ferreira et al., 2016; Ferreira et al., 2020) and herpes simplex virus 1 (Zheng and Su, 2017) have developed unique strategies to specifically target and evade the peroxisomal antiviral signalling (Ferreira et al., 2019; Ferreira et al., 2021).

Peroxisomes and mitochondria are membrane-bound and highly dynamic organelles. Besides cooperating as important antiviral platforms, they also collaborate in, among others, reactive oxygen species (ROS) metabolism and fatty acids β-oxidation (Farmer et al., 2018; Fransen et al., 2017; Islinger et al., 2018; Ribeiro et al., 2012; Schrader et al., 2020; Tilokani et al., 2018; Wanders et al., 2020). Additionally, peroxisomes and mitochondria share key components of their morphology-control machinery, such as dynamin-1-like protein (DLP1) (Koch et al., 2003; Li and Gould, 2003), mitochondrial fission factor (MFF) (Gandre-Babbe and van der Bliek, 2008; Itoyama et al., 2013) and mitochondrial fission 1 protein (FIS1) (Kobayashi et al., 2007). Mitochondrial morphology plays an important role on the MAVS-mediated antiviral response originating from this organelle (Castanier et al., 2010): upon infection, mitochondrial MAVS activation allows the induction of the mitochondrial fusion protein mitofusin-1 (MFN1), leading to the organelle’s fusion (Castanier et al., 2010; Onoguchi et al., 2010). Mitochondrial elongation/fusion is also required to enhance the interaction between MAVS and the cytosolic DNA sensing adaptor stimulator of interferon genes (STING) at the endoplasmic reticulum (ER) membranes (Castanier et al., 2010). In contrast, the relevance of peroxisome morphology for the establishment of the cellular antiviral response has not yet been established.

HCMV is a large, enveloped DNA virus belonging to the *Herpesviridae* family. HCMV infections represent one of the major causes of birth defects and opportunistic diseases in immuno-compromised patients. With a slow replication cycle, HCMV has evolved several mechanisms to evade the cellular antiviral response and cell death (Fliss and Brune, 2012; Goldmacher, 2005; Jackson et al., 2011; Marques et al., 2018). This virus encodes several immediate early proteins, such as the viral mitochondrial-localized inhibitor of apoptosis (vMIA; also known as predominant UL37 exon 1 protein (pUL37×1)) (Goldmacher et al., 1999; Ma et al., 2012). vMIA has been initially reported to localize at mitochondria and to abolish apoptosis, either by disrupting the mitochondrial transition pore formation or by blocking the permeabilization of the mitochondrial outer membrane (Goldmacher et al., 1999; McCormick et al., 2003). At mitochondria, vMIA recruits the pro-apoptotic Bcl-2 family member BAX and neutralizes it by inducing its oligomerization and membrane sequestering (Arnoult et al., 2004; Poncet et al., 2004). Through the suppression of the programmed cell death of infected cells, vMIA plays a crucial role in HCMV propagation (Arnoult et al., 2004; Goldmacher et al., 1999; Ma et al., 2012; Poncet et al., 2004; Sharon-Friling et al., 2006; Zhang et al., 2013). vMIA has also been reported to impact the modulation of the mitochondrial fission/fusion process and thus, lead to the organelle network’s disruption. While some authors have associated the perturbation of the mitochondrial network to vMIA’s anti-apoptotic function (Goldmacher, 2005), others defend that it also plays a role on the modulation of the RLR/MAVS signalling at this organelle: by inducing mitochondrial fragmentation, the contacts with the ER would be reduced, hindering the interactions between MAVS and STING (Castanier et al., 2010).

vMIA also localizes at peroxisomes, where it interacts with MAVS and inhibits the peroxisomal MAVS-dependent antiviral signalling (Magalhães et al., 2016). Peroxisomal fragmentation is also induced by vMIA but, contrarily to mitochondria, this alteration of organelle morphology does not impact the antiviral signalling inhibition (Magalhães et al., 2016).

The mechanisms by which vMIA acts towards peroxisomes, either by inhibiting antiviral signalling or disturbing organelle morphology, are still unknown. In this work, we further unravel these processes and propose a model in which vMIA inhibits antiviral signalling at peroxisomes by hindering MAVS oligomerization in an MFF-dependent manner.

Our work further demonstrates that HCMV has developed distinct mechanisms to interfere with peroxisomes and mitochondria, which may result from intrinsic differences between these two organelles and their role throughout the viral infection.

## Materials and methods

### Antibodies and plasmids

Rabbit antibodies against MFF (17090-1-AP, ProteinTech, Manchester, UK) 30, Myc-tag (71D10, 2278, Cell Signalling Technology, Beverly, MA, USA), FLAG epitope (F7425, Sigma-Aldrich, St. Louis, MO, USA), β-Actin (4967, Cell Signalling, Danvers, MA, USA), PEX14 (GTX129230, GeneTex, CA,USA) and MAVS (A300-782A, Bethyl Laboratories, TX, USA), and the mouse antibodies against MAVS (E-3, SC-166583, Santa Cruz Biotechnology, Dallas, TX, USA), p-STAT1 (Y701, BD Biosciences, San Jose, CA, USA), DLP1 (611113, BD Bioscience, San Jose, CA, USA), PMP70 (SAB4200181, Sigma-Aldrich, St. Louis, MO, USA), COXIV (4850, Cell Signalling Technology, Beverly, MA, USA) and α-Tubulin (T9026, Sigma-Aldrich, St. Louis, MO, USA) were used for immunoblotting. Mouse antibodies against PMP70 (SAB4200181, Sigma-Aldrich, St. Louis, MO, USA), Myc epitope (9E10, SC-40, Santa Cruz Biotechnology, Dallas, TX, USA) were used for immunofluorescence analyses. Additionally, mouse antibodies against Myc epitope (9E10, SC-40, Santa Cruz Biotechnology, Dallas, TX, USA) and MAVS (E-3, SC-166583, Santa Cruz Biotechnology, Dallas, TX, USA) were also used for the immunoprecipitation experiments. Species-specific anti-IgG antibodies conjugated to HRP (Bio-Rad, Richmond, CA, USA) or IRDye 800CW and IRDye 680RD secondary antibodies (LI-COR Biotechonology, Cambridge, UK) were used for immunoblotting and the fluorophores TRITC (Jackson ImmunoResearch, Cambridge, UK) and Alexa 488 (Invitrogen, Waltham, MA, USA) were used for immunofluorescence.

The plasmids GFP-RIG-I-CARD (kindly provided by Dr F. Weber, Justus-Liebig Universität Giessen, Germany) and vMIA-Myc (kindly provided by Dr V. Goldmacher, ImmunoGen Inc., Cambridge, MA, USA) were used for mammalian expression.

### Cell culture, transfections and RNA interference experiments

Mouse embryonic fibroblasts (MEFs) MAVS-PEX cells and MEFs MAVS-KO cells (described in (Dixit et al., 2010)), MEFs MAVS-MITO (described below) and human embryonic kidney (HEK) 293T cells (kindly provided by Dr M. J. Amorim, Instituto Gulbenkian para a Ciência, Portugal) were cultured in Dulbecco’s modified Eagle’s medium supplemented with 100 U/ml penicillin, 100 mg/ml streptomycin and 10% fetal bovine serum (all from GIBCO, Thermo Scientific, Waltham, MA, USA) at 37°C in a humidified atmosphere of 5% CO2. MEFs MAVS-PEX cells, MEFs MAVS-MITO and MEFs MAVS-KO cells were transfected with Lipofectamine 3000 (Invitrogen, Waltham, MA, USA) or microporated with Neon® Transfection System (Invitrogen, Waltham, MA, USA) (1700 V, width: 20, 1 pulse), following manufacturer’s instructions. HEK293T cells were incubated using 1 mg/mL of Polyethylenimine (PEI, Linear, MW 25000, Polysciences, PA, USA) at a ratio of 1:6 (DNA:PEI). Cells were fixed for organelle morphology or harvested for western blot or co-immunoprecipitation assays, 24 hrs to 72 hrs after transfection.

To knock-down the expression of MFF and DLP1 by RNA interference, 21-nucleotide small interfering RNA (siRNA) duplexes were transfected into MEFs MAVS-PEX and MEFs MAVS-MITO cells using Lipofectamine RNAiMAX (Invitrogen, Waltham, MA, USA) according to the manufacturer’s instructions. Control cells were treated with transfection mix without siRNAs complexes. Cells were assayed for silencing and organelle morphology 72h after seeding. siRNA oligonucleotides were obtained as pre-designed siRNAs as follows: MFF-sense strand: 5’-CGCUGACCUGGAACAAGGAdTdT-3’ for exon 2 30 (Ambion, Austin, TX, USA); DLP1-sense strand: 5’-UCCGUGAUGAGUAUGCUUUdTdT-3’ 31 (Ambion, Austin, TX, USA).

### Generation of stable cell lines

To generate a MEFs MAVS-MITO stable cell line, MEFs MAVS-KO were transduced with retroviruses, which were first produced by transfecting HEK293T cells with pCL-Ampho and pVSV-G (provided by Dr B. Jesus, University of Aveiro, Portugal), and MSCV2.2 IRES-GFP MAVS-MITO. Twenty-four hours upon transfection, cell media was renewed and 24 hours later cell media was collected, filtered and added to MEFs MAVS KO cells, plated 24 hours before. Transduced cells were left to grow until full confluence before being split and sorted for low GFP expression level using BD FACSARIA II.

### Viral infections

HEK293T cells were infected with SeV (Cantell strain, Charles River Laboratories, Wilmington, MA, USA) with a final concentration of 100 HA units/ml, diluted in serum and antibiotic free media. Cells were incubated for 1 hr at 37°C and, afterwards growth media containing 20% of FBS was added to cells. Infection continued for 14 hrs before cells collection.

### Immunofluorescence and microscopy

Cells grown on glass coverslips were fixed with 4% paraformaldehyde in PBS, pH 7.4, for 20 min, permeabilized with 0.2% Triton X-100 for 10 min, blocked with 1% BSA solution, for 10 min, and incubated with the indicated primary and secondary antibodies, for 1 hr at room temperature in a humid environment. Cells were then stained with Hoechst (1:2000) for 3 min, before mounting the slide. Confocal images were acquired using a Zeiss LSM 880 confocal microscope (Carl Zeiss, Oberkochen, Germany) using a Plan-Apochromat 63× and 100×/1.4 NA oil objectives, a 561 nm DPSS laser and the argon laser line 488 nm (BP 505-550 and 595-750 nm filters). Images were processed using ZEN Black and ZEN Blue software (Carl Zeiss, Oberkochen, Germany). Digital images were optimized for contrast and brightness using Adobe Photoshop (Adobe Systems, San Jose, CA, USA).

### Organelle morphology quantification

For the evaluation of organelles morphology, around six hundred cells from three independent experiments were counted for each condition, considering the size/shape and number of their peroxisomes or mitochondria. For these analyses, cells were considered as containing “fragmented organelles” when organelles were significantly smaller and in higher number than the ones from the control cells. We considered cells containing “elongated organelles” as those whose organelles had a tubular shape and were significantly longer when compared to the control cells.

### Immunoprecipitation analyses

To study the interaction between STING and vMIA, MEFs MAVS-PEX cells were co-transfected with vMIA-Myc and STING-FLAG by Lipofectamine 3000 (Invitrogen, Waltham, MA, USA). Lysates were incubated with anti-MAVS antibody for 2 hrs at 4°C on a rotary mixer. Then, 50μL of beads were added to the mixture and rotated for 10 min at room temperature. The complex was washed 3 times with PBS containing 0.1% Tween20 and then resuspended in 3x SDS-sample buffer and boiled for 10 min to elute bound proteins. Untransfected MEFs MAVS-PEX cells were used as negative control for each immunoprecipitation. In all immunoprecipitations, 50 ug of total cell lysate were used as input, and for the output the same volume of input was saved from the cell lysate extracted after incubation with the antibody and beads.

### Immunoblotting

Cells lysates were obtained by using a specific lysis buffer (25 mM Tris-HCl pH 7.5, pH 8.0, 50 mM sodium chloride, 0.5% sodium deoxycholate, 0.5% Triton X-100 and supplemented with a protease-inhibitor mix) and by passing them 20 times through a 26-gauge syringe needle. Then, samples were incubated on a rotary mixer for 30 min at 4°C, before being cleared by centrifugation (17000 x g, 15 min, 4°C). Protein concentration was determined using the Bradford assay (Bio-Rad Protein Assay, Bio-Rad, Hercules, CA, USA). Protein samples were separated by SDS-PAGE on 10% or 12.5% polyacrylamide gels, transferred to nitrocellulose (PROTAN®, Whatman®, Dassel, Germany) using a semidry apparatus or wet transfer system (Bio-Rad, Hercules, CA, USA), and analysed by immunoblotting.

Immunoblots were processed after blocking membranes with 5% milk (Molico Skimmed dry milk powder, Nestlé, Vevey, Switzerland) in TBS-T (10mM Tris pH 8, 150mM, 0.005% Tween20) and using specific primary antibodies and secondary antibodies diluted in TBS-T. Between incubations, membranes were washed 3 times for 5 min in TBS-T. Immunoblots were scanned with a Bio-Rad GS-800 calibrated imaging densitometer or ChemiDoc™ Touch Gel Imaging System (Bio-Rad, Hercules, CA, USA) for chemiluminescence detection, while Li-COR Odyssey imaging system for fluorescence detection (LI-COR Biotechonology, Cambridge, UK). Images were processed using Quantity One software (Bio-Rad, Hercules, CA, USA) or Image Studio Lite 5.2 (LI-COR Biotechonology, Cambridge, UK). Bands’ quantification was done using the volume tools from Quantity One software (Bio-Rad, Hercules, CA, USA), where the background intensity was calculated using the local background subtraction method.

### RNA extraction, cDNA synthesis and quantitative real-time polymerase chain reaction

Twenty-four hours after cells transfection, total RNA was isolated using Nzyol (NZYTech, Lisbon, PT), following manufacturer’s protocol. After quantifying RNA with DS-11 spectrophotometer (DeNovix Inc., Wilmington, DE, USA), 1 μg of total RNA was treated with 1uL DNase I (Thermo Scientific, Waltham, MA, USA). cDNA was produced from treated RNA using Revert Aid Reverse Transcriptase (Thermo Scientific, Waltham, MA, USA) and Oligo-dT15 primer (Eurofins Genomics, Ebersberg, Germany) following manufacturer’s protocol. For quantitative real-time polymerase chain reaction, 2 μL of 1:10 diluted cDNA was added to 10 μL of 2x SYBR Green qPCR Master Mix (Low Rox) (Bimake, Houston, TX, USA). The final concentration of each primer was 250 nM in 20 μL of total master mix volume. Duplicates of each sample were done, and reactions were run on 7500 Real-Time PCR System (Applied Biosystems, Waltham, MA, USA). Primer sequences were designed using Beacon Designer 7 (Premier Biosoft, Palo Alto, CA, USA) for IRF1, RSAD2 and GAPDH mouse genes. The oligonucleotides used for mouse IRF1 were 5’-GGTCAGGACTTGGATATGGAA-3’ and 5’-AGTGGTGCTATCTGGTATAATGT-3’; for mouse RSAD2 were 5’-TGTGAGCATAGTGAGCAATGG-3’ and 5’-TGTCGCAGGAGATAGCAAGA-3’; for mouse GAPDH were 5’-AGTATGTCGTGGAGTCTA-3’ and 5’-CAATCTTGAGTGAGTTGTC-3’ (Eurofins Genomics, Ebersberg, Germany) GAPDH was used as a reference gene. The thermocycling reaction was done by heating at 95°C for 3 min, followed by 40 cycles of a 12 sec denaturation step at 95°C and a 30 sec annealing/elongation step at 60 °C. The fluorescence was measured after the extension step using the Applied Biosystems software (Applied Biosystems, Waltham, MA, USA). After the thermocycling reaction, the melting step was performed with slow heating, starting at 60°C and with a rate of 1%, up to 95°C, with continuous measurement of fluorescence. Data analysis was performed using the 2−ΔΔCT method.

### Organelle-enriched fractions and sucrose gradient

HEK 293T cellular fractionation was performed by homogenizing cells in Buffer A (10mM Tris-HCl pH 7.5, 10mM KCl, 1.5mM MgCl2, 0.25M D-mannitol, supplemented with cOmplete™, EDTA-free Protease Inhibitor Cocktail (Roche, Basel, Switzerland) and passing samples gently through a 26.5-gauge syringe needle. The homogenate was cleared of nuclei and membranes by centrifugation at 1000 g for 5 min at 4°C. Mitochondria-enriched fraction pellet was obtained by centrifugation at 2000 g for 10 min at 4°C. The supernatant was then centrifuged again at 25 000 g for 25 min at 4°C to obtain the peroxisome-enriched fraction pellet. Both pellets were gently resuspended in homogenization buffer supplemented with 2% n-Dodecyl β-D-maltoside, to disrupt the organelle membrane without affecting protein’s quaternary structure.

Organelle’s fractions were then processed for the separation of MAVS oligomers by sucrose gradient (as described in (Seth et al., 2005) with minor adaptations). The organelle’s-enriched fractions were loaded in 30%-60% sucrose gradients and centrifuged at 170 000g for 2 hrs at 4°C. Starting from the top, 7 equal fractions were collected and processed for SDS-PAGE and analysed by immunoblotting.

### Statistical analysis

Statistical analysis was performed in Graph Pad Prism 9 (GraphPad Software, Inc., La Jolla, CA, USA). Data are presented as mean ± standard error mean (SEM). Differences among groups were analysed by one-way ANOVA, followed by Bonferroni’s multiple comparison test; comparisons between two groups were made using unpaired T test, and p-values of < 0.05 were considered as significant (**** - p<0.0001, *** - p<0.001, ** - p<0.01, * - p<0.05 and ns – non-significant).

## Results

### Contrarily to mitochondria, vMIA depends on the organelle fission machinery to induce peroxisome fragmentation

Although the contribution of peroxisomes, in concert with mitochondria, to the cellular antiviral response has been established (Dixit et al., 2010), the key differences between the signalling pathways, originating from these two organelles, with distinct kinetics and end products, remains unknown. HCMV vMIA has been shown to inhibit both pathways and, in parallel, induce the fragmentation of both organelles (Castanier et al., 2010; Magalhães et al., 2016). This morphology change has been pinpointed as the main trigger for the vMIA-dependent antiviral signalling inhibition at mitochondria, as it would consequently reduce its association with the ER, hindering the interaction between MAVS and STING (Castanier et al., 2010). At peroxisomes, however, the antiviral signalling inhibition induced by vMIA was shown to be independent of the organelle fragmentation (Magalhães et al., 2016).

These observed differences led us to further investigate the mechanism by which vMIA induces the morphology changes at peroxisomes, in comparison with mitochondria. To that end, we evaluated its dependence on the organelles’ fission machinery, more specifically on the cytoplasmic protein DLP1, the main responsible for the final membrane fission. vMIA’s ability to induce peroxisome fragmentation was analysed upon overexpression of Myc-tagged vMIA (vMIA-Myc) and silencing of DLP1 (via small interference RNA (siRNAs) against DLP1, siDLP1)) in mouse embryonic fibroblasts (MEFs) that contain MAVS solely at peroxisomes (MEFs MAVS-PEX (Dixit et al., 2010)) (Figure 1). Upon immunolocalization with antibodies against the Myc-tag and the peroxisomal marker PMP70, the cells were examined by confocal microscopy (Figure 1A) and the organelles’ morphological alterations were quantified (Figure 1B). As expected, silencing of DLP1 induced peroxisomal elongation when compared with control cells (Figure 1A a and b). Interestingly, vMIA was no longer able to induce peroxisome fragmentation in the absence of DLP1, as no significant differences in morphology were observed in the presence of the viral protein (Figures 1A g-i, B and S1A). We have also analysed the ability of vMIA to induce peroxisome fragmentation in the absence of MFF, another major player on peroxisomal fission and one of the main anchors of DLP1 at the organelle’s membrane. Upon analysis of MEFs MAVS PEX cells containing siMFF and vMIA-Myc (Figures 1A j-l, C and S1A), no significant changes in peroxisome morphology were detected, when compared to the elongated peroxisomes observed upon silencing of MFF (Figure 1A c, B). These results clearly show that vMIA depends on a fully functional peroxisome fission machinery to be able to induce the organelle’s fragmentation.

**Figure 1.**
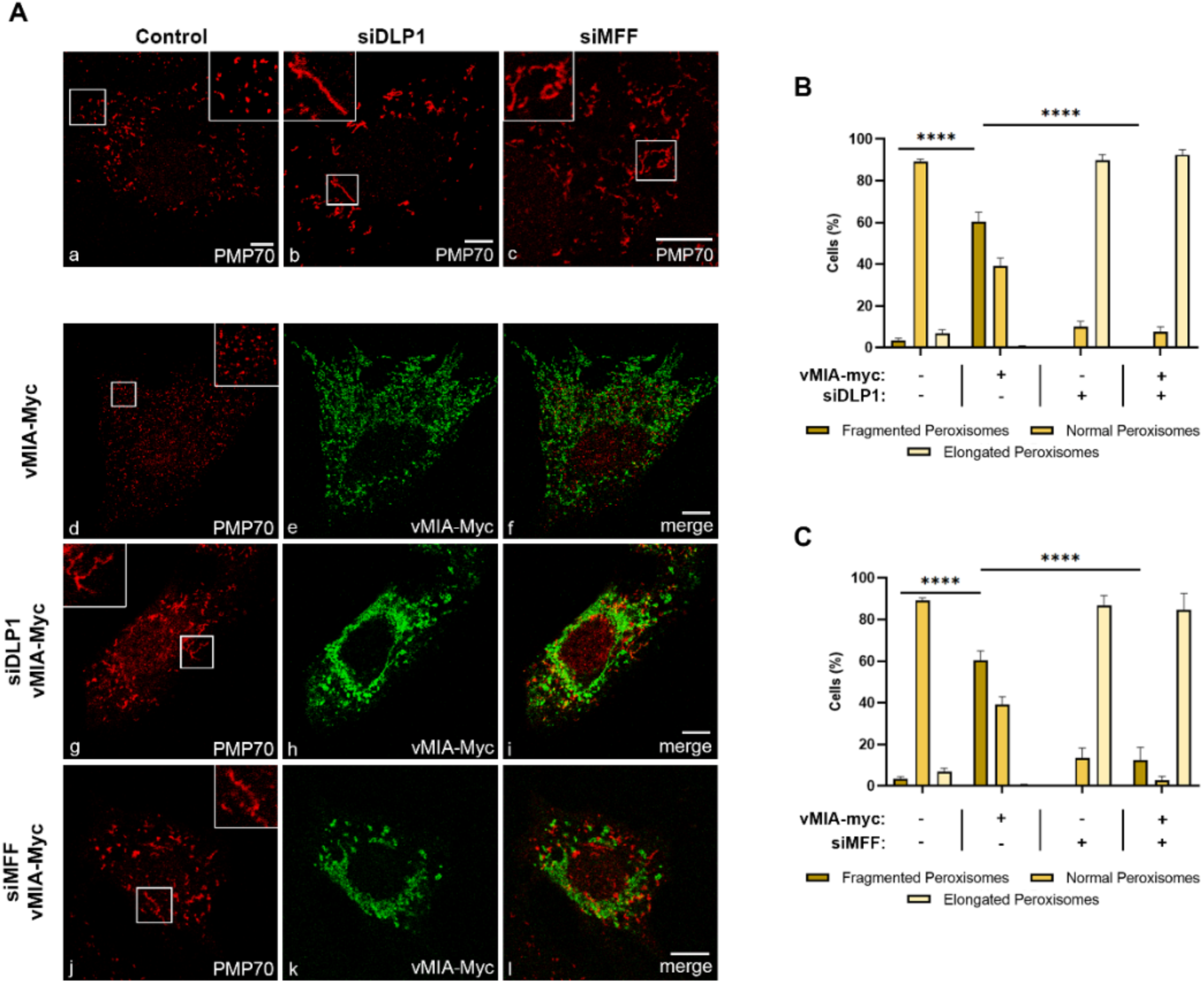
vMIA-mediated peroxisomal fragmentation is dependent on the fission machinery proteins DLP1 and MFF. (A) Immunofluorescence analyses of MEFs MAVS-PEX cells: (a) control cells, (b) DLP1 silenced cells, (c) MFF silenced cells: (a-c) anti-PMP70; (d-f) overexpression of vMIA-Myc: (d) anti-PMP70, (e) anti-Myc, (f) merge image of d and e; (g-i) overexpression of vMIA-Myc in DLP1 silenced cells: (g) anti-PM70, (h) anti-Myc, (i) merge image of g and h; (j-l) overexpression of vMIA-Myc in MFF silenced cells: (j) anti-PMP70, (k) anti-Myc, (l) merge image of j and k. Bars represent 10 μm. (B and C) Statistical analysis of peroxisomal morphology upon overexpression of vMIA-Myc in MEFs MAVS-PEX cells in the absence of DLP1 and MFF, respectively. Approximately 600 cells were analysed per condition. Data represents the means ± SEM of three independent experiments analysed using two-way ANOVA with Bonferroni’s multi comparations test (ns = non-significant, **** - p < 0.0001). Error bars represent SEM.

In order to analyse whether an analogous dependence occurs at mitochondria, we performed similar analyses in MEFs MAVS-MITO cells (where MAVS is present solely at mitochondria). Mitochondria morphology was analysed by confocal microscopy upon immunolocalization with antibodies against Myc and the mitochondrial marker TIM23, in cells expressing vMIA-Myc in the presence or absence of DLP1 or MFF (siDLP1 or siMFF) (Figure 2). Surprisingly, vMIA was still able to induce mitochondria fragmentation in the absence of either DLP1 or MFF (Figure S1B), totally reverting the organelle elongation observed upon silencing of any of these proteins (Figure 2A g-l, B and C). These results were also observed upon analysis of mitochondria morphology in MEFs MAVS PEX cells, in the absence of DLP1 or MFF (Figure S1C g-l, D and E). This data indicates that, contrarily to peroxisomes, vMIA is able to interfere with mitochondria morphology in a fission machinery-independent manner.

**Figure 2.**
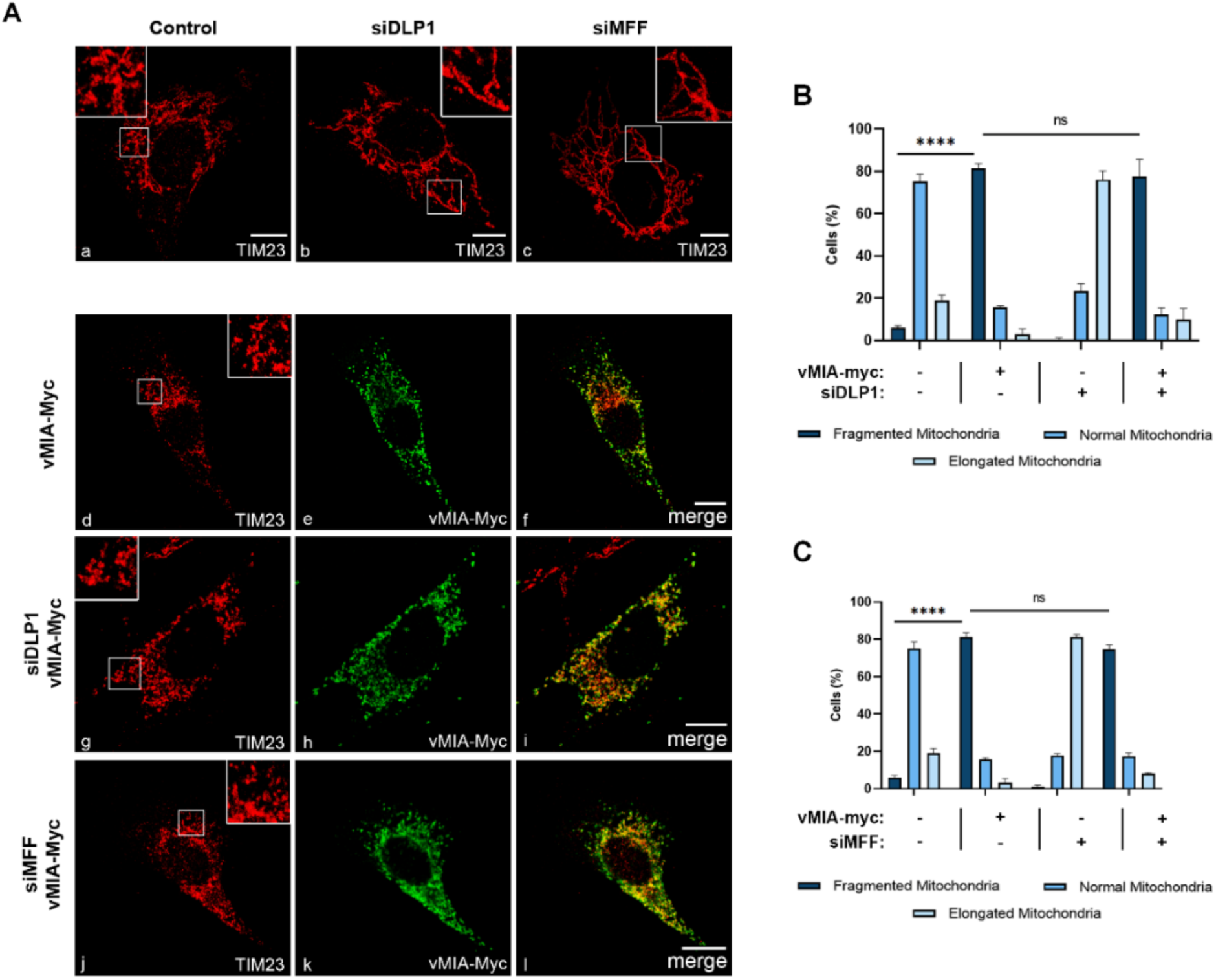
vMIA-mediated mitochondrial fragmentation is independent on the fission machinery proteins DLP1 and MFF. (A) Immunofluorescence analyses of MEFs MAVS-MITO cells: (a) control cells, (b) DLP1 silenced cells, (c) MFF silenced cells: (a-c) anti-TIM23; (d-f) overexpression of vMIA-Myc: (d) anti-TIM23, (e) anti-Myc, (f) merge image of d and e; (g-i) overexpression of vMIA-Myc in DLP1 silenced cells: (g) anti-TIM23, (h) anti-Myc, (i) merge image of g and h; (j-l) overexpression of vMIA-Myc in MFF silenced cells: (j) anti-TIM23, (k) anti-Myc, (l) merge image of j and k. Bars represent 10 μm. (B and C) Statistical analysis of mitochondrial morphologies upon overexpression of vMIA-Myc in MEFs MAVS-MITO cells in the absence of DLP1 and MFF, respectively. Approximately 600 cells were analysed per condition. Data represents the means ± SEM of three independent experiments analysed using two-way ANOVA with Bonferroni’s multi comparations test (ns = non-significant, **** - p < 0.0001). Error bars represent SEM.

### vMIA-induced peroxisome and mitochondria fragmentation is independent of MAVS signalling

The independence on the organelle morphology for the vMIA-induced antiviral signalling inhibition at peroxisomes (Magalhães et al., 2016), contrarily to what was observed at mitochondria (Castanier et al., 2010), suggests that these two processes may be less correlated than initially anticipated. In order to conclude on whether the peroxisomal and mitochondrial fragmentation prompted by vMIA is in any way dependent on MAVS signalling, we analysed the organelles’ morphology in cells lacking MAVS, in the presence of the viral protein. To that end, MEFs MAVS-KO cells were transfected with vMIA-Myc, and confocal microscopy analyses were performed upon immunolocalization with antibodies against PMP70 and TIM23. As shown in Figure 3A, even in the absence of MAVS, vMIA was able to induce a strong fragmentation of both peroxisomes (Figure 3B) and mitochondria (Figure 3C). These results demonstrate that the organelle morphology and the mechanism of vMIA-mediated inhibition of the MAVS signalling are less related than anticipated.

**Figure 3.**
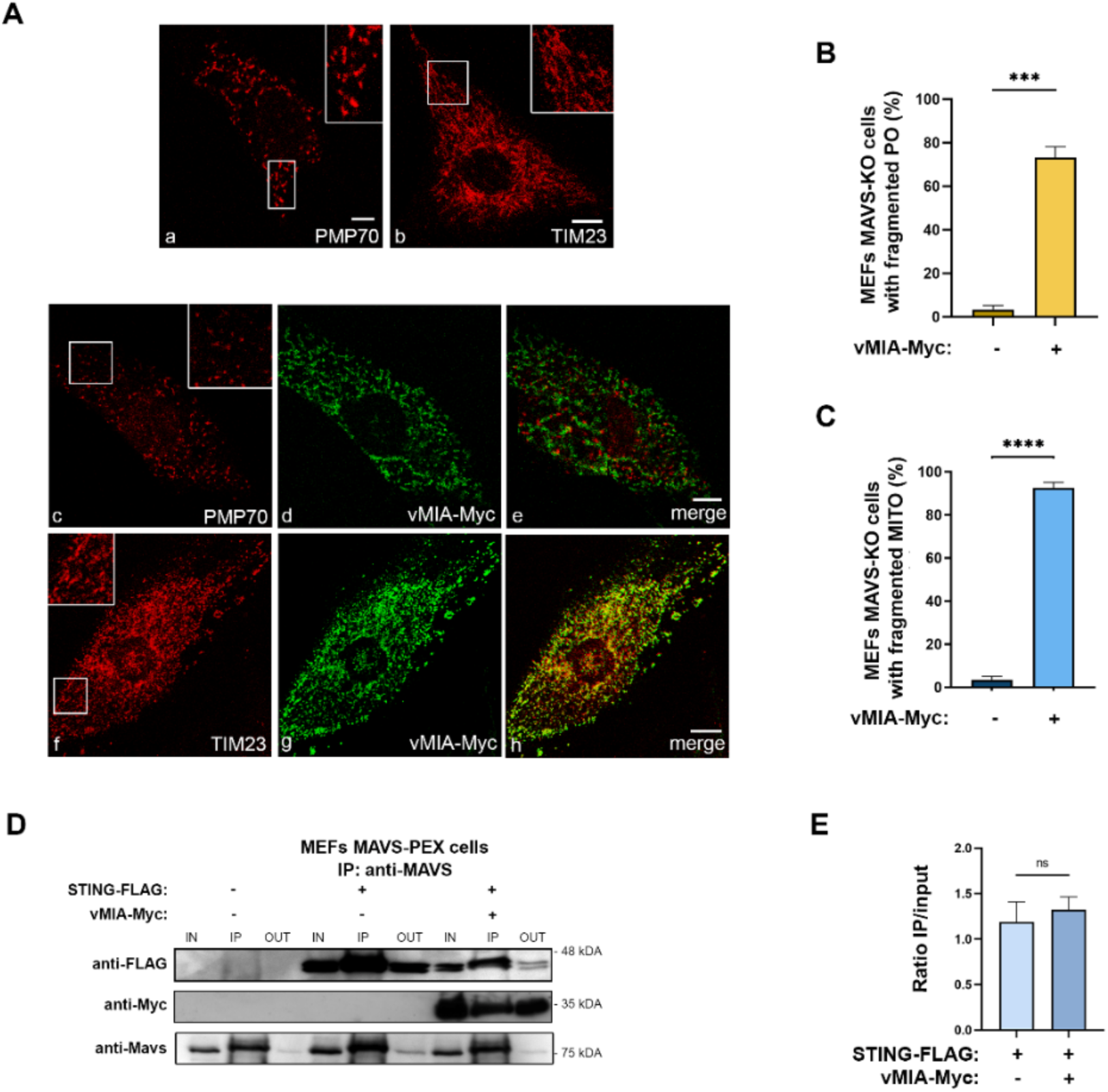
vMIA-induced peroxisomal and mitochondrial fragmentation is independent of MAVS. vMIA does not disrupt STING-MAVS interaction at peroxisomes. (A) Immunofluorescence analyses of MEFs MAVS KO cells: (a-b) peroxisomal and mitochondrial morphologies in control cells: (a) anti-PMP70, (b) anti-TIM23; (c-e) peroxisomal morphology upon overexpression of vMIA-Myc: (c) anti-PMP70, (d) anti-Myc, (e) merge image of c and d; (f-h) mitochondrial morphology upon overexpression of vMIA-Myc: (f) anti-TIM23, (g) anti-Myc, (h) merge image of f and g. Bars represent 10 μm. (B and C) Statistical analysis of peroxisomal and mitochondrial morphologies upon overexpression of vMIA-Myc in MEFs MAVS KO cells, respectively. Approximately 600 cells were analysed per condition. (D) Co-immunoprecipitation analysis of the interaction between overexpressed STING-FLAG and vMIA-Myc in MEFs MAVS-PEX cells. The pull-down was performed using an antibody against MAVS. Western blot was performed with antibodies against FLAG and Myc. IN represents total cell lysate (input), IP represents immunoprecipitation and OUT represents the cell lysate extracted after incubation with the antibody (output). (E) Quantification of the ratio between IP and IN, in the presence or absence of vMIA. Data represents the means ± SEM of three independent experiments, analysed using unpaired T test (ns - non-significant; *** - p<0.001, **** - p<0.0001).

Nevertheless, as the mitochondria fragmentation induced by vMIA has been associated to a decrease in MAVS-STING interaction and the consequent inhibition of the immune response (Castanier et al., 2010), we analysed whether the same would be observed upon peroxisomal fragmentation. We started by analysing the occurrence of an interaction between the peroxisomal MAVS and STING, as this had not yet been demonstrated. To that end, we transfected MEFs MAVS-PEX with a FLAG-tagged STING (STING-FLAG) and, 24 hrs after, performed co-immunoprecipitation analyses with a pull-down against MAVS. As shown in Figure 3D, STING interacts with the peroxisomal MAVS. In order to investigate whether this interaction is eventually compromised by vMIA, these experiments were also performed upon co-transfection with vMIA-Myc. As shown in Figure 3D and 3E, the presence of the viral protein does not inhibit the interaction between STING and peroxisomal MAVS, contrarily to what had been observed at mitochondria. These results, together with the distinct dependencies on organelle morphology, point out major differences between the mechanisms involved in the modulation of the peroxisomal and mitochondrial antiviral signalling by vMIA.

### MFF is essential for the vMIA-mediated inhibition of the peroxisome-dependent antiviral signalling

To further unravel the mechanism by which vMIA inhibits the MAVS signalling at peroxisomes, we once more evaluated the importance of the fission machinery proteins DLP1 and MFF. Upon silencing of any of these proteins in MEFs MAVS-PEX cells, vMIA-Myc was transfected and, 24 hrs after, the MAVS-dependent antiviral signalling was stimulated by overexpressing a constitutively active form of RIG-I (GFP-RIG-I-CARD, as in (Magalhães et al., 2016; Yoneyama et al., 2004)). In correlation with our previous results, the absence of DLP1 did not hinder the peroxisomal antiviral response, leading to similar expression levels of ISGs (IRF1 and RSAD2) when comparing with the ones obtained in the stimulated control cells (Figure 4). However, our results clearly show that, at peroxisomes, MFF’s absence strongly impairs the vMIA-mediated inhibition of IRF1 and RSAD2 mRNA expression (Figure 4A). MFF, hence, rises as the first identified peroxisomal protein to play an important role on the mechanism of vMIA-mediated inhibition of the peroxisome-dependent antiviral response.

**Figure 4.**
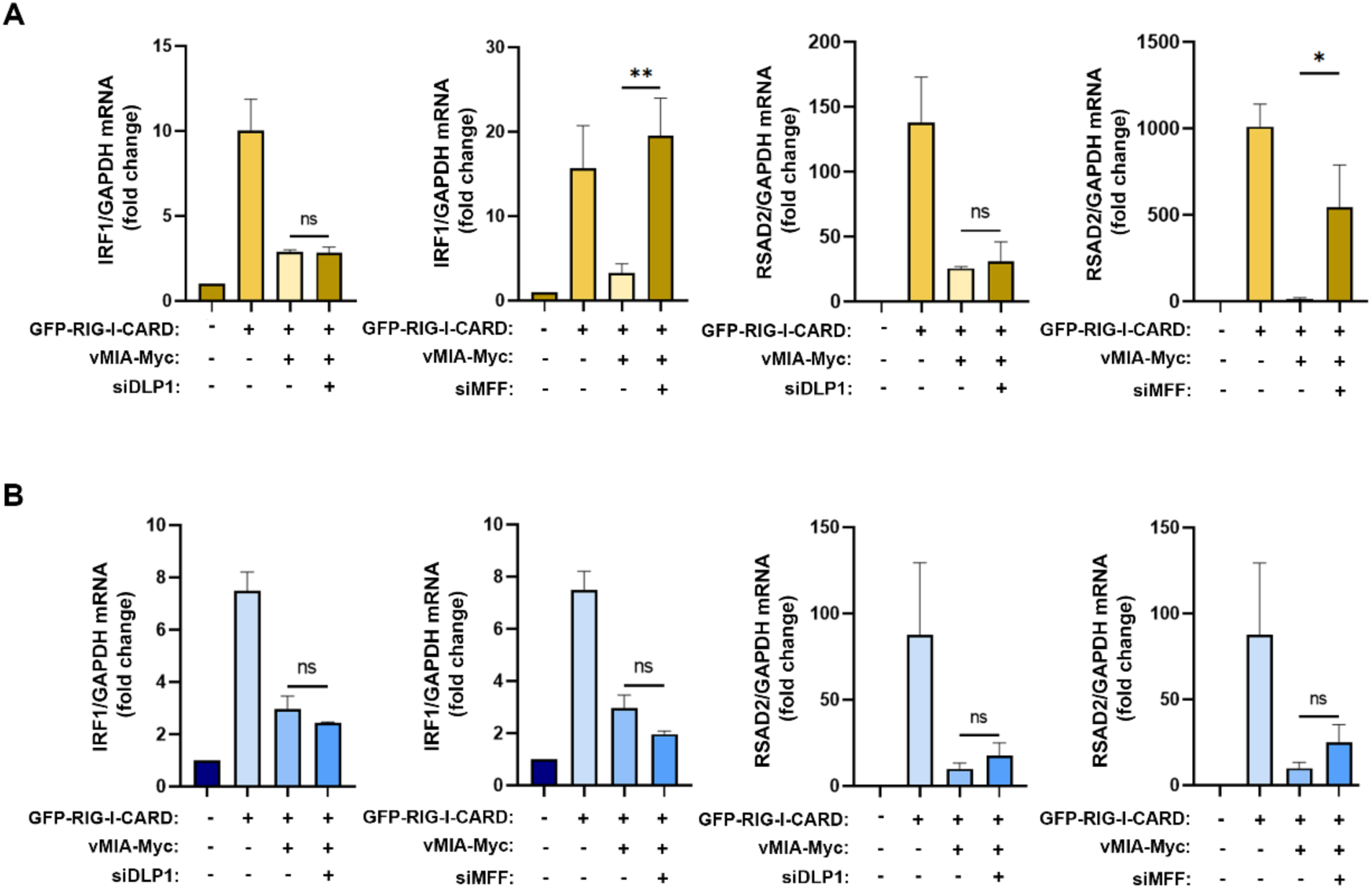
MFF is required for vMIA-mediated inhibition of the MAVS signalling at peroxisomes but not at mitochondria. (A and B) RT-qPCR analysis of IRF1 and RSAD2 (viperin) mRNA expression in (A) MEFs MAVS-PEX cells or (B) MEFs MAVS-MITO cells stimulated with GFP-RIG-I-CARD, upon vMIA-Myc overexpression in the absence of DLP1 or MFF. GAPDH was used as normalizer gene and graphs depict fold change in comparation to untreated samples. Data represents the means ± SEM of at least three independent experiments analysed using one-way ANOVA with Bonferroni’s multi comparations test (ns - non-significant; * - p<0.005; ** - p<0.01).

In order to determine whether the same dependency is observed at mitochondria, similar experiments were performed in MEFs MAVS-MITO cells. Surprisingly, in these cells, neither the lack of DLP1 nor MFF had significant effects on ISGs expression (Figure 4B).

The importance of MFF for the antiviral signalling inhibition mediated by vMIA seems to be, hence, exclusive for the peroxisomal response.

### vMIA inhibits MAVS oligomerization at peroxisomes and mitochondria

One of the crucial steps of the cellular RIG-I/MAVS antiviral signalling, is the oligomerization of MAVS at the peroxisomal and mitochondrial membranes upon interaction with RIG-I, forming functional high molecular weight prion-like aggregates, essential for the activation of the downstream proteins (Hou et al., 2011; Seth et al., 2005). To further unravel the mechanism of action of vMIA towards the peroxisomal MAVS signalling, and as we had previously observed that this viral protein interacts with MAVS, we analysed its effect on MAVS oligomerization upon infection with Sendai virus (SeV) in HEK293T cells. As these cells contain MAVS at both peroxisomes and mitochondria, we used differential centrifugation to prepare peroxisome-enriched fractions and analyse solely the oligomerization of peroxisomal MAVS. Gradient assay experiments were then implemented to separate peroxisomal MAVS in different density fractions, depending on its oligomerization degree. As shown in Figure 5A, upon infection with SeV, confirmed by the increase in p-STAT1 (Figure 5E), peroxisomal MAVS oligomers appear at the higher density fraction. Interestingly, this is no longer observed in the presence of vMIA (Figure 5A). These results suggest that this viral protein, likely through direct interaction (previously reported in (Magalhães et al., 2016)), prevents the formation of MAVS oligomers, inhibiting the downstream antiviral signalling (Figure 5A and E).

**Figure 5.**
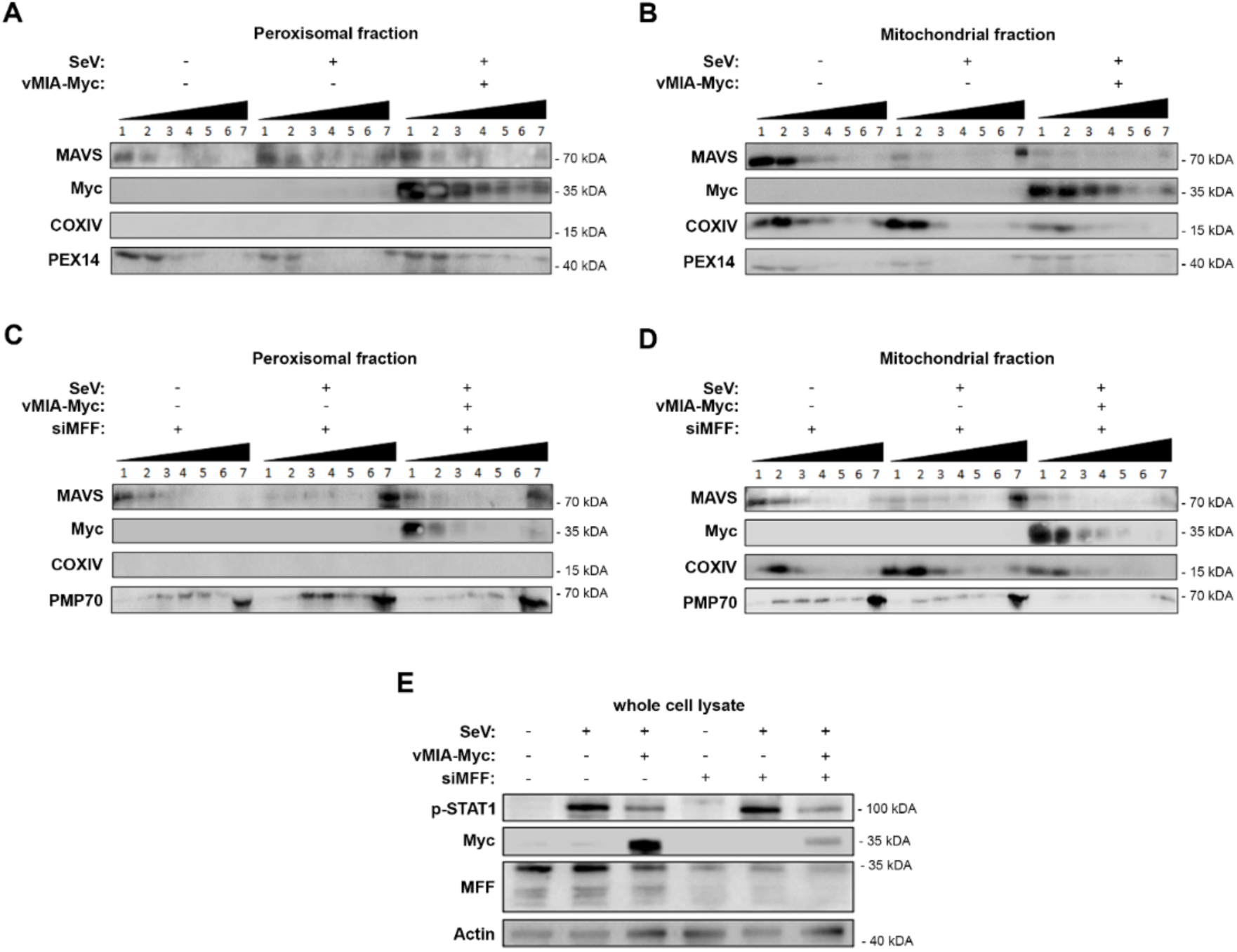
vMIA inhibits MAVS oligomerization at peroxisomes and mitochondria. MFF is essential for the vMIA-mediated inhibition of MAVS oligomerization at peroxisomes but not at mitochondria. (A and B) HEK293T cells infected with SeV in the presence or absence of vMIA. Density gradient assay was performed to demonstrate the separation of endogenous MAVS based on its density. 1 – 7 represent the fractions isolated from the gradient assay, where 1 represents the fraction with lowest density and 7 represents the fraction with highest density. (A) Peroxisome-enriched fraction, (B) Mitochondria-enriched fraction. (C and D) HEK293T cells infected with SeV in the presence or absence of vMIA and in the absence of MFF. Density gradient assay was performed to demonstrate the separation of endogenous MAVS based on its density. 1 – 7 represent the fractions isolated from the gradient assay, where 1 represents the fraction with lowest density and 7 represents the fraction with highest density. (C) Peroxisome-enriched fraction, (D) Mitochondria-enriched fraction. (A, B, C and D) Immunoblots were performed with antibodies against MAVS, Myc-tag, COXIV, PEX14 and PMP70. (E) Whole cell lysates were resolved by SDS-PAGE. SeV infection, and consequential activation of MAVS downstream signalling, was confirmed using anti-p-STAT1. vMIA-Myc overexpression and MFF silencing were also confirmed using anti-Myc and anti-MFF, respectively. Antibody against Actin was used as loading control.

In order to unravel whether this oligomerization inhibition also occurs at the mitochondrial membranes, we have performed similar analysis on mitochondria-enriched fractions isolated by differential centrifugation. As shown in Figure 5B, the results were in all similar to the ones obtained in peroxisomes, demonstrating that, vMIA is also able to inhibit mitochondrial MAVS oligomerization and, consequently, the downstream antiviral signalling (Figure 5B and E).

### vMIA-mediated inhibition of MAVS oligomerization at peroxisomes, but not at mitochondria, is dependent on MFF

The dependency on MFF for the vMIA influence towards the peroxisomal antiviral response, led us to investigate whether this protein would be involved in the observed inhibition of MAVS oligomerization. Similar gradient separation analysis of peroxisome-enriched fractions was performed in MFF silenced (via siMFF transfection) HEK293T cells infected with SeV. Interestingly, our results show that, in the absence of MFF, vMIA is no longer capable of inhibiting the formation of peroxisomal MAVS oligomers, and hence the downstream antiviral signalling (Figure 5C, E).

On the other hand, similar experiments performed in mitochondrial-enriched fractions of HEK293T cells infected with SeV, indicate that vMIA is still able to inhibit MAVS oligomerization in the absence of MFF and, hence, hinder the downstream antiviral signalling (Figure 5 D and E). These results correlate to our previous observation that MFF is not essential for the vMIA-mediated inhibition of the antiviral signalling at mitochondria.

Altogether our results demonstrate that vMIA inhibits MAVS oligomerization at peroxisomes in an MFF-dependent manner, and highlight important differences between its mechanisms of action towards both organelles.

## Discussion

The significance of peroxisomes for the establishment of the cellular immune response has been highlighted by different reports demonstrating that distinct viruses have developed specific strategies to target and evade the peroxisomal antiviral signalling (Berg et al., 2012; Cohen et al., 2000; Ferreira et al., 2016; Ferreira et al., 2019; Ferreira et al., 2021; Han et al., 2014; Jefferson et al., 2014; Xu et al., 2017; You et al., 2015; Zheng and Su, 2017). We have also previously demonstrated that HCMV interferes with the peroxisomal antiviral pathway through its protein vMIA (Magalhães et al., 2016). In that study, we show that vMIA interacts with PEX19 to be transported to the peroxisomal membranes, where it interacts with MAVS, induces the organelle’s fragmentation, and inhibits the antiviral immune response. Peroxisomal fragmentation has, however, been shown not to influence the vMIA-induced inhibition of the antiviral signalling, suggesting that two vMIA-mediated independent processes are occurring at this organelle (Magalhães et al., 2016). These results have also uncovered dissimilarities between the mechanisms of action of vMIA towards peroxisomes and mitochondria, as, at mitochondria, vMIA inhibition of the antiviral signalling is dependent on the organelle’s fission (Castanier et al., 2010). These observations were not initially anticipated, as these organelles, besides sharing MAVS, are dependent on the same fission machinery, harbouring proteins, such as MFF (Gandre-Babbe and van der Bliek, 2008), which anchors DLP1 at the membranes of both organelles for final fission (Imoto et al., 2020).

The reasoning behind the vMIA-mediated inhibition of the antiviral response at mitochondria has been suggested to be based on the disruption of the MAVS-STING interactions due to the reduction of mitochondria-ER contact sites, as a consequence of the organelle’s fragmentation (Castanier et al., 2010). Here, we show, for the first time, that STING and MAVS also interact at the peroxisomal level but, contrarily to mitochondria, this interaction is not disturbed by the presence of vMIA. This result uncovers key differences between the mechanisms of action of vMIA at these two organelles, which may reflect intrinsic, yet unidentified, dissimilarities between the two antiviral signalling pathways.

By evaluating vMIA’s dependence on the organelles’ fission machineries to fragment peroxisomes and mitochondria, we have unraveled further discrepancies between the two pathways. While at peroxisomes vMIA is not able to induce fragmentation in the absence of DLP1 and MFF, these proteins are not essential for its action towards mitochondrial fragmentation. It is then tempting to suggest that, at peroxisomes, vMIA somehow stimulates MFF to recruit more DLP1 and induce organelle fragmentation. At mitochondria, however, the organelle fragmentation induced by vMIA may be related to the role of this viral protein on the control of apoptosis, interfering with BAX to prevent mitochondrial outer-membrane permeabilization, and mediating the release of ER Ca^2+^ stores into the cytosol (Ma et al., 2012; Poncet et al., 2006; Sharon-Friling et al., 2006). One could hypothesize that one of vMIA’s roles at peroxisomes could also be related with its anti-apoptotic function. Although a direct influence of peroxisomes on apoptosis has not yet been demonstrated, the anti-apoptotic proteins Bcl2 and Bcl-XL, as well as the pro-apoptotic protein BAK (Costello et al., 2017; Fujiki et al., 2017; Hosoi et al., 2017) have also been identified at peroxisomes. It has also been demonstrated that peroxisomal BAK regulates membrane integrity and the release of soluble peroxisomal matrix proteins, such as catalase (Fujiki et al., 2017; Hosoi et al., 2017). The presence of anti-apoptotic proteins, such as vMIA, at the peroxisomal membranes could, hence, protect the organelle from excessive matrix protein release into the cytosol. However, it has been shown that, at least in mitochondria, vMIA does not block BAK-mediated apoptosis and that this inhibition is mediated by another HCMV protein (Arnoult et al., 2004; Cam et al., 2010).

Besides showing that the antiviral signalling inhibition is unrelated to the organelle morphology changes, we have also demonstrated that the vMIA-induced peroxisomal fragmentation is totally independent of the presence of MAVS at the organelle’s membranes. It was recently shown that HCMV induces the upregulation of peroxisomal proteins and peroxisome growth and fission, mainly at late times post-infection, increasing peroxisome numbers and leading to a higher production of plasmalogens, contributing to the enhancement of virus production (Beltran et al., 2018). In a subsequent study, the authors suggest a model by which vMIA activates PEX11β, which in turn induces MFF to upregulate peroxisome fission during infection (Federspiel et al., 2020). These results are in line with our data and together indicate that vMIA induces peroxisome fragmentation to favor viral propagation at late times post-infection, while, earlier in infection, it specifically inhibits the peroxisome-dependent signalling.

Our results clearly also demonstrate that MFF is essential for the vMIA-mediated inhibition of the immune response at peroxisomes, but not at mitochondria, revealing once more important differences between the mechanisms occurring at both organelles. Importantly, we further reveal that vMIA, likely though direct interaction (Magalhães et al., 2016), inhibits MAVS oligomerization at both peroxisomes and mitochondria, further impairing the downstream signalling. Coherently, MFF has been shown to be essential for this oligomerization inhibition at peroxisomes, but not at mitochondria.

Based on all our results, we suggest a model for vMIA’s mechanism of action towards peroxisomes, which is depicted in Figure 6: upon infection, vMIA interacts with PEX19 at the cytoplasm and travels to peroxisomes, where it interacts with MAVS. This interaction interferes, in an MFF-dependent manner, with the formation of MAVS oligomers and inhibits the consequent activation of the downstream antiviral signalling. In parallel, vMIA induces peroxisome fragmentation in a MFF- and DLP1-dependent manner, but independently of MAVS, which may be important for the enhancement of lipid metabolism and virus particles formation at late times post-infection.

**Figure 6.**
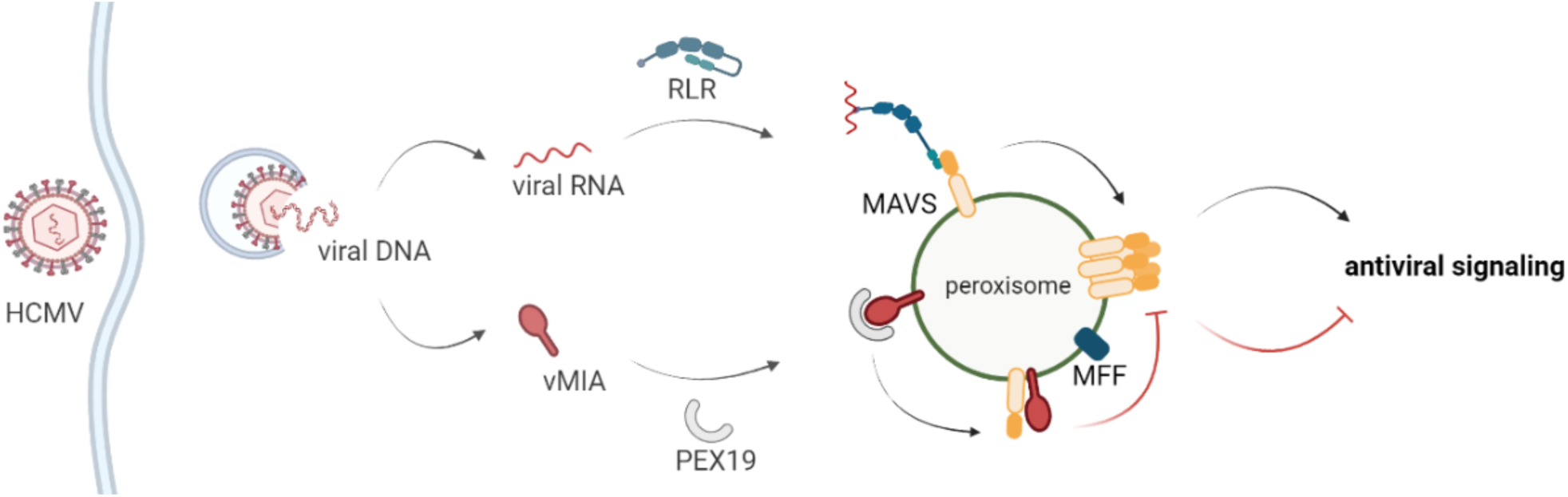
Schematic representation of the proposed model for the vMIA-dependent inhibition of the peroxisomal antiviral signalling. Upon stimulation, RIG-I-like proteins interact with MAVS at peroxisomes, inducing its oligomerization and the downstream production of direct antiviral effectors. HCMV vMIA is able to specifically evade this peroxisomal antiviral signalling. vMIA interacts with PEX19 at the cytoplasm and travels to peroxisomes, where it interacts with MAVS. This interaction interferes, in an MFF-dependent manner, with the formation of MAVS oligomers and inhibits the consequent activation of the downstream antiviral signalling.

In conclusion, in this manuscript we not only propose the molecular mechanism by which HCMV evades the peroxisomal antiviral response, but also shed some light on possible molecular processes that may be occurring at mitochondria. Our results once more emphasize the relevance of peroxisomes as platforms for antiviral signalling against HCMV and uncover molecular mechanisms that may be explored as targets for antiviral therapy.

## Author Contributions

AF, AG, JK and DR contributed to conception and design of the study. AF, AG, AM, IV and MM acquired the data; AF, AG, JK and DR wrote the manuscript; All authors contributed to manuscript revision, read, and approved the submitted version.

## Acknowledgments

We thank Dr Victor Goldmacher for kindly providing the vMIA-Myc plasmid, Dr Friedemann Weber for kindly providing the GFP-RIG-I-CARD plasmid, Dr Maria João Amorim for providing the HEK293T cells and Dr Bruno Jesus for providing pCL-Ampho and pVSV-G. The authors thank Dr Michael Schrader, Dr Markus Islinger and Dr Jorge Azevedo for the valuable discussions. The authors also thank all the members of the Virus Host-cell Interactions Laboratory (especially Vanessa Ferreira and Bruno Ramos, who prepared the MEFs MAVS-MITO cell line) and Kagan’s Laboratory, for the valuable inputs and discussions. Image acquisition was performed in the LiM facility of iBiMED, a node of PPBI (Portuguese Platform of BioImaging): POCI-01-0145-FEDER-022122.

## References

Arnoult, D., Bartle, L. M., Skaletskaya, A., Poncet, D., Zamzami, N., Park, P. U., Sharpe, J., Youle, R. J. and Goldmacher, V. S. (2004). Cytomegalovirus cell death suppressor vMIA blocks Bax-but not Bak-mediated apoptosis by binding and sequestering Bax at mitochondria. Proc. Natl. Acad. Sci. U. S. A. 101, 7988–93.

Beltran, P. M. J., Cook, K. C., Hashimoto, Y., Galitzine, C., Murray, L. A., Vitek, O. and Cristea, I. M. (2018). Infection-Induced Peroxisome Biogenesis Is a Metabolic Strategy for Herpesvirus Replication. Cell Host Microbe 24, 1–16.

Bender, S., Reuter, A., Eberle, F., Einhorn, E., Binder, M. and Bartenschlager, R. (2015). Activation of Type I and III Interferon Response by Mitochondrial and Peroxisomal MAVS and Inhibition by Hepatitis C Virus. PLOS Pathog. 11, e1005264.

Berg, R. K., Melchjorsen, J., Rintahaka, J., Diget, E., Søby, S., Horan, K. A., Gorelick, R. J., Matikainen, S., Larsen, C. S., Ostergaard, L., et al. (2012). Genomic HIV RNA induces innate immune responses through RIG-I-dependent sensing of secondary-structured RNA. PLoS One 7, e29291.

Cam, M., Handke, W., Picard-Maureau, M. and Brune, W. (2010). Cytomegaloviruses inhibit Bak-and Bax-mediated apoptosis with two separate viral proteins. Cell Death Differ. 17, 655–665.

Castanier, C., Garcin, D., Vazquez, A., Arnoult, D., Ablasser, A., Bauernfeind, F., Hartmann, G., Latz, E., Fitzgerald, K., Hornung, V., et al. (2010). Mitochondrial dynamics regulate the RIG-I-like receptor antiviral pathway. EMBO Rep. 11, 133–138.

Cohen, G. B., Rangan, V. S., Chen, B. K., Smith, S. and Baltimore, D. (2000). The human thioesterase II protein binds to a site on HIV-1 Nef critical for CD4 down-regulation. J. Biol. Chem. 275, 23097–23105.

Costello, J. L., Castro, I. G., Camões, F., Schrader, T. A., McNeall, D., Yang, J., Giannopoulou, E.-A., Gomes, S., Pogenberg, V., Bonekamp, N. A., et al. (2017). Predicting the targeting of tail-anchored proteins to subcellular compartments in mammalian cells. J. Cell Sci. 130, 1675–1687.

Dixit, E. and Kagan, J. C. (2013). Intracellular Pathogen Detection by RIG-I-Like Receptors. Adv. Immunol. 117, 99–125.

Dixit, E., Boulant, S., Zhang, Y., Lee, A. S. Y., Odendall, C., Shum, B., Hacohen, N., Chen, Z. J., Whelan, S. P., Fransen, M., et al. (2010). Peroxisomes are signalling platforms for antiviral innate immunity. Cell 141, 668–81.

Farmer, T., Naslavsky, N. and Caplan, S. (2018). Tying Trafficking to Fusion and Fission at the Mighty Mitochondria. Traffic 19, 569–577.

Federspiel, J. D., Cook, K. C., Kennedy, M. A., Venkatesh, S. S., Otter, C. J., Hofstadter, W. A., Jean Beltran, P. M. and Cristea, I. M. (2020). Mitochondria and Peroxisome Remodeling across Cytomegalovirus Infection Time Viewed through the Lens of Inter-ViSTA. Cell Rep. 32, 107943.

Ferreira, A. R., Magalhães, A. C., Camões, F., Gouveia, A., Vieira, M., Kagan, J. C. and Ribeiro, D. (2016). Hepatitis C virus NS3-4A inhibits the peroxisomal MAVS-dependent antiviral signalling response. J. Cell. Mol. Med. 20, 750–757.

Ferreira, A. R., Marques, M. and Ribeiro, D. (2019). Peroxisomes and Innate Immunity: Antiviral Response and Beyond. Int. J. Mol. Sci. 20, 3795.

Ferreira, A. R., Ramos, B., Nunes, A. and Ribeiro, D. (2020). Hepatitis C Virus: Evading the Intracellular Innate Immunity. J. Clin. Med. 9, 790.

Ferreira, A. R., Marques, M., Ramos, B., Kagan, J. C. and Ribeiro, D. (2021). Emerging roles of peroxisomes in viral infections. Trends Cell Biol. 0,.

Fliss, P. M. and Brune, W. (2012). Prevention of cellular suicide by cytomegaloviruses. Viruses 4, 1928–49.

Fransen, M., Lismont, C. and Walton, P. (2017). The peroxisome-mitochondria connection: How and why? Int. J. Mol. Sci. 18, 1126.

Fujiki, Y., Miyata, N., Mukai, S., Okumoto, K. and Cheng, E. H. (2017). BAK regulates catalase release from peroxisomes. Mol. Cell. Oncol. 4, e1306610.

Gandre-Babbe, S. and van der Bliek, A. M. (2008). The novel tail-anchored membrane protein Mff controls mitochondrial and peroxisomal fission in mammalian cells. Mol. Biol. Cell 19, 2402–12.

Goldmacher, V. S. (2005). Cell death suppression by cytomegaloviruses. Apoptosis 10, 251–265.

Goldmacher, V. S., Bartle, L. M., Skaletskaya, A., Dionne, C. A., Kedersha, N. L., Vater, C. A., Han, J. W., Lutz, R. J., Watanabe, S., Cahir McFarland, E. D., et al. (1999). A cytomegalovirus-encoded mitochondria-localized inhibitor of apoptosis structurally unrelated to Bcl-2. Proc. Natl. Acad. Sci. U. S. A. 96, 12536–41.

Han, J.-M., Kang, J.-A., Han, M.-H., Chung, K.-H., Lee, C.-R., Song, W.-K., Jun, Y. and Park, S.-G. (2014). Peroxisome-localized hepatitis Bx protein increases the invasion property of hepatocellular carcinoma cells. Arch. Virol. 159, 2549–57.

Horner, S. M., Liu, H. M., Park, H. S., Briley, J. and Gale, M. (2011). Mitochondial-associated endoplasmic reticulum membranes (MAM) form innate immune synapses and are targeted by hepatitis C virus. Proc Natl Acad Sci 108, 14590–14595.

Hosoi, K.-I., Miyata, N., Mukai, S., Furuki, S., Okumoto, K., Cheng, E. H. and Fujiki, Y. (2017). The VDAC2-BAK axis regulates peroxisomal membrane permeability. J. Cell Biol. 216, 709–722.

Hou, F., Sun, L., Zheng, H., Skaug, B., Jiang, Q.-X. and Chen, Z. J. (2011). MAVS forms functional prionlike aggregates to activate and propagate antiviral innate immune response. Cell 146, 448–461.

Imoto, Y., Itoh, K. and Fujiki, Y. (2020). Molecular Basis of Mitochondrial and Peroxisomal Division Machineries. Int. J. Mol. Sci. 21, 5452.

Islinger, M., Voelkl, A., Fahimi, H. D. and Schrader, M. (2018). The peroxisome: an update on mysteries 2.0. Histochem. Cell Biol. 150, 443–471.

Itoyama, A., Michiyuki, S., Honsho, M., Yamamoto, T., Moser, A., Yoshida, Y. and Fujiki, Y. (2013). Mff functions with Pex11p and DLP1 in peroxisomal fission. Biol. Open 2, 998–1006.

Jackson, S. E., Mason, G. M. and Wills, M. R. (2011). Human cytomegalovirus immunity and immune evasion. Virus Res. 157, 151–160.

Jefferson, M., Whelband, M., Mohorianu, I. and Powell, P. P. (2014). The pestivirus N terminal protease Npro redistributes to mitochondria and peroxisomes suggesting new sites for regulation of IRF3 by Npro. PLoS One 9, e88838.

Kawai, T., Takahashi, K., Sato, S., Coban, C., Kumar, H., Kato, H., Ishii, K. J., Takeuchi, O. and Akira, S. (2005). IPS-1, an adaptor triggering RIG-I-and Mda5-mediated type I interferon induction. Nat. Immunol. 6, 981–988.

Kell, A. M. and Gale, M. (2015). RIG-I in RNA virus recognition. Virology 0, 110–121.

Kobayashi, S., Tanaka, A. and Fujiki, Y. (2007). Fis1, DLP1, and Pex11p coordinately regulate peroxisome morphogenesis. Exp. Cell Res. 313, 1675–1686.

Koch, A., Thiemann, M., Grabenbauer, M., Yoon, Y., McNiven, M. a and Schrader, M. (2003). Dynamin-like protein 1 is involved in peroxisomal fission. J. Biol. Chem. 278, 8597–605.

Li, X. and Gould, S. J. (2003). The dynamin-like GTPase DLP1 is essential for peroxisome division and is recruited to peroxisomes in part by PEX11. J. Biol. Chem. 278, 17012–20.

Ma, J., Edlich, F., Bermejo, G. a, Norris, K. L., Youle, R. J. and Tjandra, N. (2012). Structural mechanism of Bax inhibition by cytomegalovirus protein vMIA. Proc. Natl. Acad. Sci. U. S. A. 109, 20901–6.

Magalhães, A. C., Ferreira, A. R., Gomes, S., Vieira, M., Gouveia, A., Valença, I., Islinger, M., Nascimento, R., Schrader, M., Kagan, J. C., et al. (2016). Peroxisomes are platforms for cytomegalovirus’ evasion from the cellular immune response. Sci. Rep. 6, 26028.

Marques, M., Ferreira, A. R. and Ribeiro, D. (2018). The Interplay between Human Cytomegalovirus and Pathogen Recognition Receptor Signaling. Viruses 10, 514.

McCormick, A. L., Smith, V. L., Chow, D. and Mocarski, E. S. (2003). Disruption of Mitochondrial Networks by the Human Cytomegalovirus UL37 Gene Product Viral Mitochondrion-Localized Inhibitor of Apoptosis. J. Virolo 77, 631–641.

Meylan, E., Curran, J., Hofmann, K., Moradpour, D., Binder, M., Bartenschlager, R. and Tschopp, J. (2005). Cardif is an adaptor protein in the RIG-I antiviral pathway and is targeted by hepatitis C virus. Nature 437, 1167–72.

Onoguchi, K., Onomoto, K., Takamatsu, S., Jogi, M., Takemura, A., Morimoto, S., Julkunen, I., Namiki, H., Yoneyama, M. and Fujita, T. (2010). Virus-infection or 5’ppp-RNA activates antiviral signal through redistribution of IPS-1 mediated by MFN1. PLoS Pathog. 6, e1001012.

Poncet, D., Larochette, N., Pauleau, A. L., Boya, P., Jalil, A. A., Cartron, P. F., Vallette, F., Schnebelen, C., Bartle, L. M., Skaletskaya, A., et al. (2004). An anti-apoptotic viral protein that recruits Bax to mitochondria. J. Biol. Chem. 279, 22605–22614.

Poncet, D., Pauleau, A.-L., Szabadkai, G., Vozza, A., Scholz, S. R., Le Bras, M., Brière, J.-J., Jalil, A., Le Moigne, R., Brenner, C., et al. (2006). Cytopathic effects of the cytomegalovirus-encoded apoptosis inhibitory protein vMIA. J. Cell Biol. 174, 985–996.

Ribeiro, D., Castro, I., Fahimi, H. D. and Schrader, M. (2012). Peroxisome morphology in pathology. Histol. Histopathol. 27, 661–76.

Saito, T. and Gale, M. (2008). Differential recognition of double-stranded RNA by RIG-I-like receptors in antiviral immunity. J. Exp. Med. 205, 1523–7.

Schrader, M., Kamoshita, M. and Islinger, M. (2020). Organelle interplay—peroxisome interactions in health and disease. J. Inherit. Metab. Dis. 43, 71–89.

Seth, R. B., Sun, L., Ea, C.-K. and Chen, Z. J. (2005). Identification and Characterization of MAVS, a Mitochondrial Antiviral Signaling Protein that Activates NF-κB and IRF3. Cell 122, 669–682.

Sharon-Friling, R., Goodhouse, J., Colberg-Poley, A. M. and Shenk, T. (2006). Human cytomegalovirus pUL37×1 induces the release of endoplasmic reticulum calcium stores. Proc. Natl. Acad. Sci. U. S. A. 103, 19117–22.

Tilokani, L., Nagashima, S., Paupe, V. and Prudent, J. (2018). Mitochondrial dynamics: overview of molecular mechanisms. Essays Biochem. 62, 341–360.

Wanders, R. J. A., Vaz, F. M., Waterham, H. R. and Ferdinandusse, S. (2020). Fatty Acid Oxidation in Peroxisomes: Enzymology, Metabolic Crosstalk with Other Organelles and Peroxisomal Disorders. Adv. Exp. Med. Biol. 1299, 55–70.

Xu, L.-G., Wang, Y.-Y., Han, K.-J., Li, L.-Y., Zhai, Z. and Shu, H.-B. (2005). VISA is an adapter protein required for virus-triggered IFN-beta signaling. Mol. Cell 19, 727–40.

Xu, Z., Asahchop, E. L., Branton, W. G., Gelman, B. B., Power, C. and Hobman, T. C. (2017). MicroRNAs upregulated during HIV infection target peroxisome biogenesis factors: Implications for virus biology, disease mechanisms and neuropathology. PLoS Pathog. 13, e1006360.

Yoneyama, M., Kikuchi, M., Natsukawa, T., Shinobu, N., Imaizumi, T., Miyagishi, M., Taira, K., Akira, S. and Fujita, T. (2004). The RNA helicase RIG-I has an essential function in double-stranded RNA-induced innate antiviral responses. Nat. Immunol. 5, 730–7.

You, J., Hou, S., Malik-Soni, N., Xu, Z., Kumar, A., Rachubinski, R. A., Frappier, L. and Hobman, T. C. (2015). Flavivirus infection impairs peroxisome biogenesis and early anti-viral signaling. J. Virol. 89, 12349–61.

Zhang, A., Hildreth, R. L. and Colberg-Poley, A. M. (2013). Human cytomegalovirus inhibits apoptosis by proteasome-mediated degradation of bax at endoplasmic reticulum-mitochondrion contacts. J. Virol. 87, 5657–68.

Zheng, C. and Su, C. (2017). Herpes simplex virus 1 infection dampens the immediate early antiviral innate immunity signaling from peroxisomes by tegument protein VP16. Virol. J. 14, 35.

