## Supplementary Figure 1 for "Human cytomegalovirus vMIA inhibits MAVS oligomerization at peroxisomes in an MFF-dependent manner"

### Supplementary Material

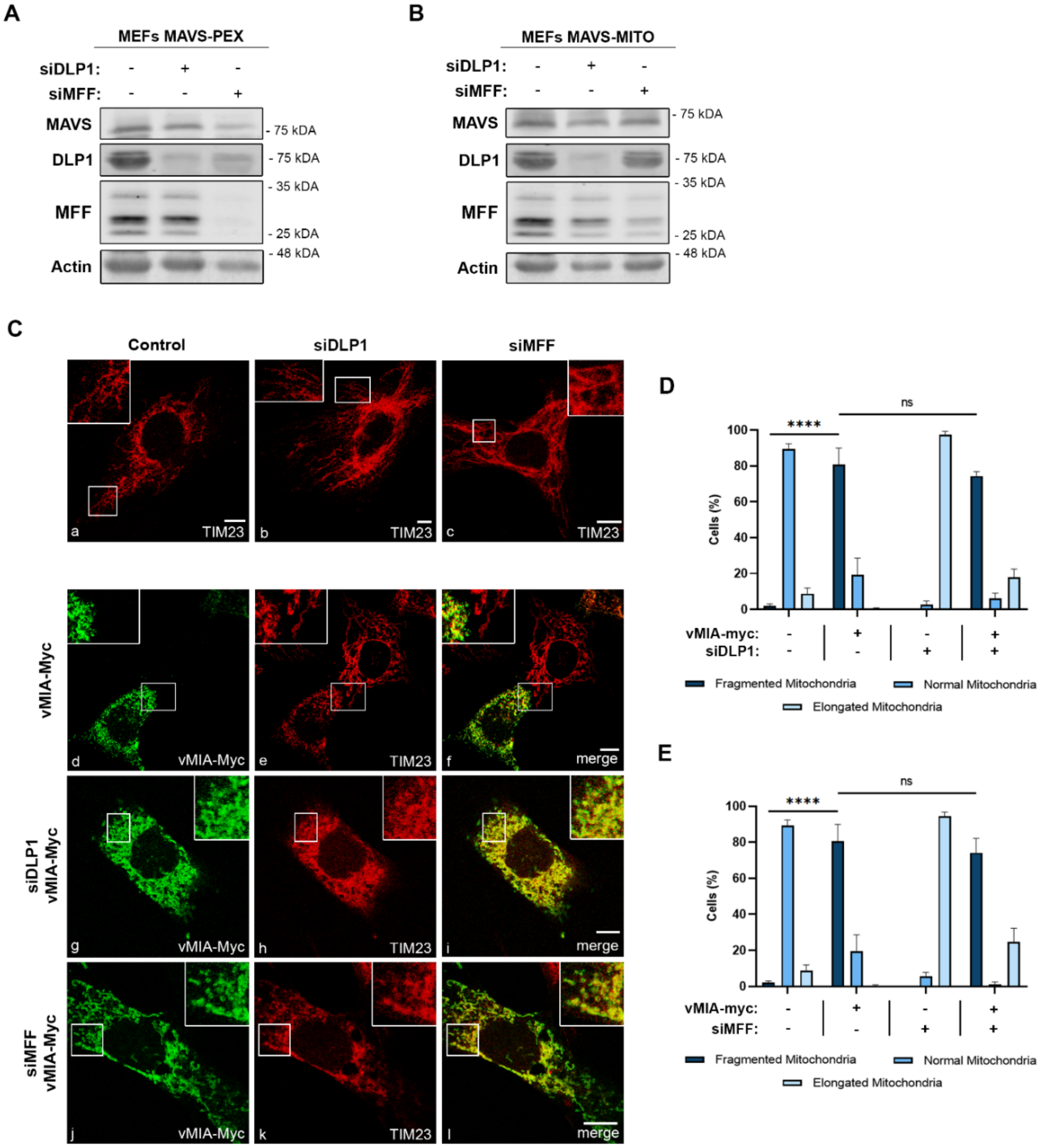


**Supplementary Figure 1.** (A) Western blot analysis of MEFs MAVS-PEX cells to confirm DLP1 and MFF silencing in the experiments represented in Figure 1. (B) Western blot analysis of MEFs MAVS-MITO cells to confirm DLP1 and MFF silencing in the experiments represented in Figure 2. (A and B) Immunoblots were performed with antibodies against MAVS, DLP1 and MFF. Actin was used as loading control. (C) Immunofluorescence analyses of MEFs MAVS-PEX cells: (a) control cells, (b) DLP1 silenced cells, (c) MFF silenced cells: (a-c) anti-TIM23; (d-f) overexpression of vMIA-Myc: (d) anti-TIM23, (e) anti-Myc, (f) merge image of d and e; (g-i) overexpression of vMIA-Myc in DLP1 silenced cells: (g) anti-TIM23, (h) anti-Myc, (i) merge image of g and h; (j-l) overexpression of vMIA-Myc in MFF silenced cells: (j) anti-TIM23, (k) anti-Myc, (l) merge image of j and k. Bars represent 10 µm. (D and E) Statistical analysis of mitochondrial morphologies upon overexpression of vMIA-Myc in MEFs MAVS-PEX cells in the absence of DLP1 and MFF, respectively. Approximately 600 cells were analyzed per condition. Data represents the means ± SEM of three independent experiments analyzed using two-way ANOVA with Bonferroni's multi comparations test (ns = non-significant, **** - $p < 0.0001$). Error bars represent SEM.
